# Direct conversion of human fibroblasts to pancreatic epithelial cells through transient progenitor states is controlled by temporal activation of defined factors

**DOI:** 10.1101/2022.11.16.516750

**Authors:** Liangru Fei, Kaiyang Zhang, Nikita Poddar, Sampsa Hautaniemi, Biswajyoti Sahu

## Abstract

Cell fate can be reprogrammed by ectopic expression of lineage-specific transcription factors (TF). For example, few specialized cell types like neurons, hepatocytes and cardiomyocytes have been generated from fibroblasts by defined factors (Wang *et al*, 2021). However, the exact cell state transitions and their control mechanisms during cell fate conversion are still poorly understood. Moreover, the defined TFs for generating vast majority of the human cell types are still elusive. Here, we report a novel protocol for reprogramming human fibroblasts to pancreatic exocrine cells with phenotypic and functional characteristics of ductal epithelial cells using a minimal set of six TFs. We mapped the molecular determinants of lineage dynamics at single-cell resolution using a novel factor-indexing method based on single-nuclei multiome sequencing (FI-snMultiome-seq) that enables dissecting the role of each individual TF and pool of TFs in cell fate conversion. We show that transdifferentiation – although being considered a direct cell fate conversion method – occurs through transient progenitor states orchestrated by stepwise activation of distinct TFs. Specifically, transition from mesenchymal fibroblast identity to epithelial pancreatic exocrine fate involves two deterministic steps: first, an endodermal progenitor state defined by activation of HHEX concurrently with FOXA2 and SOX17, and second, temporal GATA4 activation essential for maintenance of pancreatic cell fate program. Collectively, our data provide a high-resolution temporal map of the epigenome and transcriptome remodeling events that facilitate cell fate conversion, suggesting that direct transdifferentiation process occurs through transient dedifferentiation to progenitor cell states controlled by defined TFs.

## Main

Direct cell fate conversion (transdifferentiation) is a process in which somatic cells are reprogrammed to defined cell types of another lineage without a pluripotent intermediate state using either ectopic expression of cell- and lineage-specific TFs, non-coding RNAs or small molecules in a highly defined lineage-dependent media (Wang *et al*, 2021). For example, fibroblasts have been directly converted to cells representing all three germ layers, including induced neuronal cells from the ectoderm (Vierbuchen *et al*, 2010), cardiomyocytes from the mesoderm (Ieda *et al*, 2010) and hepatocytes from the endoderm (Sekiya *et al*, 2011). Transdifferentiation is controlled by coordinated action of pioneer TFs such as FOXA or GATA factors and other lineage-specific TFs, resulting in global reprogramming of epigenetic landscape and gene expression (Iwafuchi-Doi *et al*, 2014). Previous studies have demonstrated considerable cellular heterogeneity during lineage conversion (Biddy *et al*, 2018, Francesconi *et al*, 2019, Schiebinger *et al*, 2019), but the exact path of cellular states through which the direct lineage conversion occurs is not fully understood.

Transdifferentiation approaches have been critical not only for understanding basic developmental mechanisms governing cell identity, but also for innovative experimental strategies in disease modelling and potential therapeutic applications. These include, for example, conversion of astrocytes to dopaminergic neurons in Parkinson’s disease (Qian *et al*, 2020) and glial cells into neurons after brain injury (Guo *et al*, 2014), as well as combining direct transdifferentiation from fibroblasts to induced hepatocytes with controlled expression of cancer-specific oncogenes to model liver cancer development (Sahu *et al*, 2021). However, the lack of transdifferentiation factors for many human cell and tissue types hampers the use of this approach for understanding various human diseases. Pancreas is a complex organ that harbors multiple specialized cell types with endocrine (α, β, δ, γ, and ε cells) and exocrine (acinar and ductal cells) functions. Enormous efforts have gone into identifying the TFs required for generation of endocrine β-cells (Zhou *et al*, 2008, Lee *et al*, 2013, Chen *et al*, 2014, Ariyachet *et al*, 2016) but the factors required for other defined cell types are currently unknown. However, the exocrine cells such as the ductal epithelial cells are important not only for the normal physiological function of the pancreas but also in various disease contexts (Ellis *et al*, 2017). For example, chronic pancreatitis and cystic fibrosis impair exocrine function, and highly lethal pancreatic cancer that is often diagnosed at the terminal stages typically originates from the exocrine cells. Recent functional studies have implicated the lineage-determining TFs in tumorigenic processes (Sahu *et al*, 2021, Patel *et al*, 2022, Baggiolini *et al*, 2021). Thus, it is pertinent to study the role of defined TFs in controlling pancreatic exocrine cell identity for better understanding of pancreatic cancer as well as for potential regenerative medicine applications.

Here, we report the defined factors necessary and sufficient for direct conversion of human fibroblasts to induced pancreatic exocrine cells (iPEC) of ductal epithelial identity. We have dissected the mechanistic role of individual TFs in the transdifferentiation process, revealing the critical factors that control the process in a coordinated spatio-temporal manner.

### Generation of pancreatic exocrine cells by direct lineage conversion

To establish a direct cell fate conversion protocol for transdifferentiating human fibroblasts to iPECs in defined media (**Fig. 1a**), we set out to identify the pool of TFs required for inducing pancreatic exocrine cell fate. We selected a total of 14 candidate TFs (**Supplementary Fig. 1a**), eight of which were literature-curated based on their reported role in maturation of exocrine cells during human pancreas development (Jennings *et al*, 2015, Petersen *et al*, 2018), and another eight predicted using the Mogrify computational framework (Rackham *et al*, 2016). Two of the TFs, FOXA2 and GATA4, were suggested by both approaches. The candidate TFs were cloned into a lentiviral expression vector and systematically studied in a series of reprogramming experiments by transducing 20 different TF combinations to human foreskin fibroblasts (HFF; **Supplementary Fig. 1b**). Transduced cells were monitored for morphological changes and for the expression of pancreas-related marker genes using quantitative RT-PCR (qRT-PCR) or bulk RNA-sequencing (RNA-seq) at different time points (**Supplementary Fig. 1b;** see **Methods** for details). Briefly, after observing clear morphological changes in a pilot experiment (condition 1; **Supplementary Fig. 1b, c**), we compared the gene expression profiles of transduced cells to control fibroblasts and detected upregulation of pancreatic exocrine cell markers as well as enrichment of pancreas-related gene set among the differentially expresses genes (DEG) (**Supplementary Fig. 1d-f**). We then designed two sub-pools for generating acinar- and ductal-like cells separately (conditions 2 and 3 with ten and nine TFs, respectively; **Supplementary Fig. 1b**). Reprogramming experiments using these pools demonstrated the feasibility of producing iPECs, since qRT-PCR and RNA-seq analyses showed a gradual increase in the expression of marker genes for acinar and ductal cells and downregulation of fibroblast-related genes (**Supplementary Fig. 1g-j**). The pools were further refined based on the previous literature about pancreas development (Petersen *et al*, 2018, Villamayor *et al*, 2020, Solar *et al*, 2009, Schaffer *et al*, 2010, Shih *et al*, 2012, Hale *et al*, 2014), resulting in eight and six TFs for acinar and ductal cells, respectively (conditions 5 and 7; **Supplementary Fig. 1b**; see **Methods** for details). The cells transduced with acinar cell TFs did not develop epithelial-like morphology. However, the cells transduced with a pool of six TFs for ductal cells, FOXA2, SOX17, PDX1, HNF1B, HNF6 (ONECUT1) and SOX9 (henceforth referred to as 6F), showed clear morphological changes during transdifferentiation with several clusters of epithelial-like cells appearing around three weeks of reprogramming (**Supplementary Fig. 2a**). Dissociating the cells and re-plating them on growth factor reduced Matrigel-coated dishes resulted in further maturation towards epithelial cell-like morphology – cells of the epithelial-like clusters that were picked and re-plated at four weeks after induction could be expanded up to eight weeks as the cells mature and differentiate (**Fig. 1b**). To facilitate molecular characterization of the reprogrammed cells at later timepoints, these cells were immortalized by lentiviral expression of hTERT.

**Figure 1.**
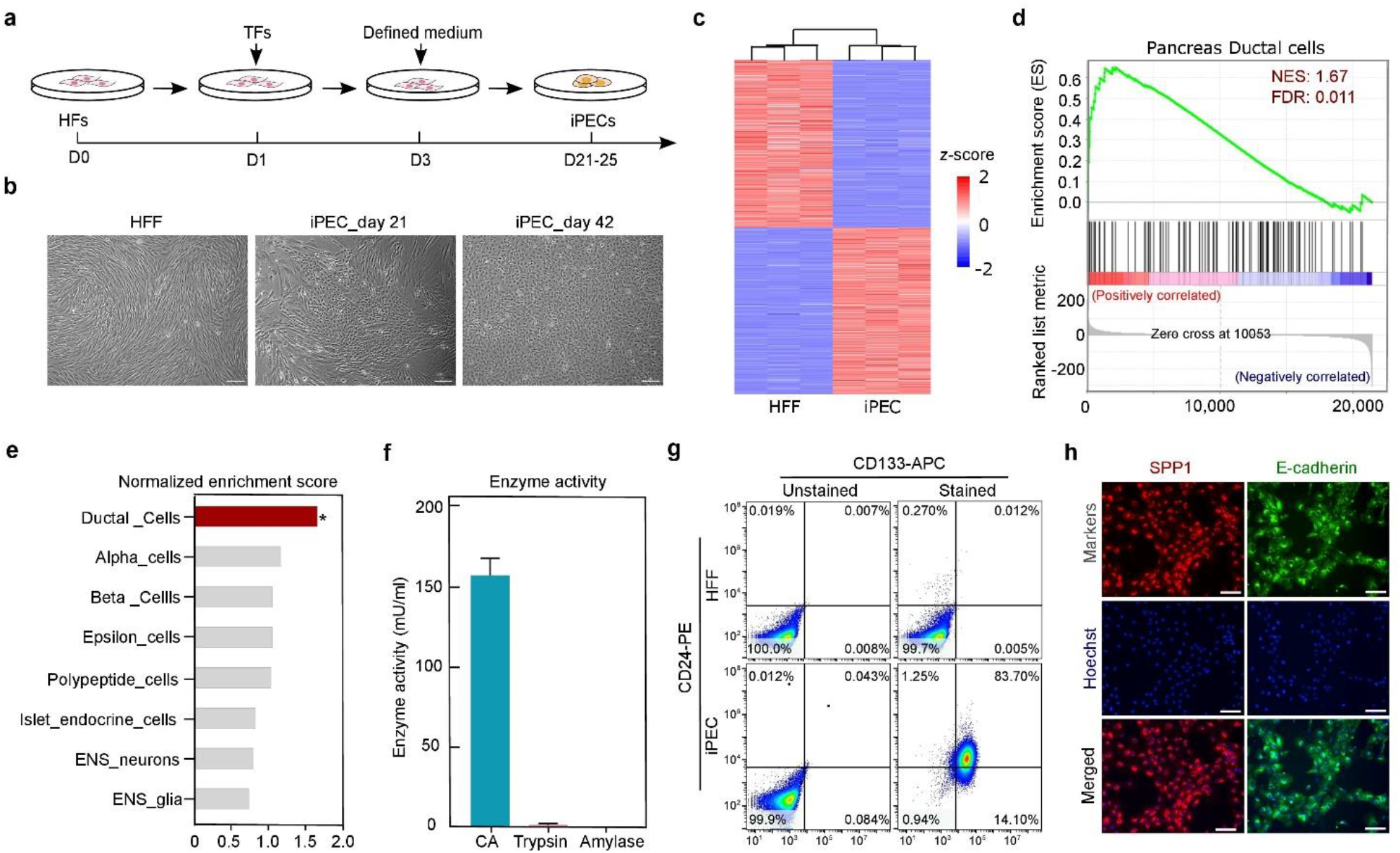
Direct conversion of human fibroblasts to pancreatic ductal-like cells. **a**, Experimental design to test candidate TFs required for generating iPECs. Human foreskin fibroblasts (HFF) were transduced with lentiviral expression constructs for different combinations of candidate TFs. The culture medium was changed to defined reprogramming medium two days after transduction (see **Methods**), and the cells were monitored for morphological changes. Reprogrammed iPECs display epithelial-like morphology 21-25 days after 6F transduction. **b**, Bright field images showing spindle-shaped HFFs and epithelial-like morphology of iPECs at day 21 and 42 after induction. **c**, Hierarchical clustering of differentially expressed genes (DEG) in HFFs vs. iPECs at 10wk^TERT^ (|log2FC| ≥ 1.5, *p*-value < 0.05). *z*-score of normalized expression values is indicated using a color scale. **d**, Gene set enrichment analysis (GSEA) showing significantly enriched fetal pancreatic ductal cell signatures in iPECs at 10wk^TERT^. **e**, Bar plot showing normalized enrichment scores from GSEA for signatures of different pancreatic cell types within upregulated genes in iPECs at 10wk^TERT^ with specific enrichment for ductal cell signature (*, *p*-value < 0.01 and false discovery rate (FDR) < 5%). **f**, Measurement of enzyme activities for CA, trypsin and amylase in the lysates of iPECs at eight weeks after induction. Data are represented as the mean ± SEM (n = 6). **g**, Flow cytometry analysis for the expression of CD133 and CD24 to determine the proportion of ductal-like cells in the iPEC population at six weeks after induction. HFFs were used as control. Of note, CD133 protein is encoded by the *PROM1* gene (see also **Supplementary Fig. 1h**). **h**, Immunofluorescence staining for SPP1, E-cadherin and cellular DNA (Hoechst) to assess the epithelial phenotype of iPECs at six weeks after induction. (Scale bars in all cell images throughout this manuscript represent 100 μm).

To characterize the identity of cells generated using the 6F combination, RNA-seq was performed from reprogrammed cells at six and ten weeks after TF transduction (using TERT-immortalized cells for the later timepoint, 10wk^TERT^). Global gene expression analysis revealed a distinct transcriptional program within the reprogrammed cells compared to HFFs (**Fig.1c**). Pancreatic marker genes such as secreted phosphoprotein 1 (*SPP1*) and carbonic anhydrase 2 (*CA2*) were upregulated and the genes highly expressed in mesenchymal-origin fibroblast cells were efficiently downregulated (**Supplementary Fig. 2b**), which is in agreement with earlier reports of hepatic, cardiac and neuronal conversion from fibroblasts (Ieda *et al*, 2010, Huang *et al*, 2014, Treutlein *et al*, 2016). Importantly, gene set enrichment analysis (GSEA) for DEGs between 6F-reprogrammed cells and control fibroblasts showed statistically significant positive enrichment specifically for pancreatic ductal cell gene signature among signatures for all major cell types of endodermal origin (**Supplementary Fig. 2c, d**). Furthermore, among all pancreatic cell types, only the ductal cell signature was significantly enriched in 10wk^TERT^ cells (**Fig. 1d, e**). These results indicate that the 6F combination induces cell fate conversion towards iPECs with a pancreatic ductal cell identity.

### Cell fate conversion results in iPECs with functional properties of pancreatic ductal cells

For testing the functional properties of iPECs induced with 6F, we performed enzyme activity assays for three pancreatic enzymes: carbonic anhydrase (CA) that is the key enzyme expressed in pancreatic ducts, as well as amylase and trypsin that are specific for acinar cells. Strong CA activity was detected in iPECs (**Fig. 1f**), whereas amylase and trypsin activities were negligible. These results confirm that the cells transduced with 6F pool are of pancreatic ductal cell identity.

To evaluate the proportion of ductal cells in the isolated iPEC colonies, we performed flow cytometry analysis using a well-known ductal cell surface marker CD133, and CD24 that has been shown to be restricted to exocrine cells of human pancreas (Muraro *et al*, 2016). Over 80% of iPECs expressed both CD133 and CD24, demonstrating high purity of ductal cells in the isolated iPEC colonies (**Fig. 1g**). Moreover, clear immunofluorescence (Hale *et al*) signal was detected for SPP1, an essential regulator of human pancreatic duct cell maturation and gatekeeper of epithelial phenotype of ductal cells (Hendley *et al*, 2021), and for epithelial marker E-cadherin (**Fig. 1h**), further demonstrating the ductal phenotype of iPECs and efficient conversion of mesenchymal cells to epithelial cells. Taken together, the 6F combination of TFs can efficiently and directly convert human fibroblasts towards pancreatic epithelial cells of ductal identity with functional characteristics of ductal epithelial cells.

### iPEC reprogramming is controlled by temporal activation of specific TFs

To gain insights into the transcriptional changes during direct lineage conversion between two somatic cell fates – fibroblasts and pancreatic ductal cells – the cells were collected for bulk RNA-seq analysis at different timepoints during transdifferentiation. Transcriptome-wide changes were analyzed using principal component analysis (PCA) (**Supplementary Fig. 2e**), and pair-wise differential gene expression analysis was performed for reprogrammed cells at each timepoint against HFF control. Unsupervised hierarchical clustering for the significant DEGs overlapping all pair-wise comparisons revealed two major clusters (**Fig. 2a**), the fibroblast-related genes that were silenced or down-regulated during reprogramming and the pancreas-related genes that showed dynamic gene expression changes during transdifferentiation process. Gene Ontology (GO) terms related to epithelial cell differentiation were significantly enriched among the genes that were rapidly induced within the first week of reprogramming (**Fig. 2a**). This is in line with recent studies showing that mesenchymal-epithelial transition is an essential early step in somatic cell reprogramming (Pei *et al*, 2019). Rapid downregulation of fibroblast-related genes already at 48 h timepoint and strong induction of ductal cell marker genes such as *SALL4, ACSM3, HABP2, CA2* and *SPP1* during later stages of reprogramming (**Fig. 2a**) indicate a successful cell fate switch between fibroblasts and iPECs and gradual maturation of the ductal cells. In agreement with this, pancreatic ductal cell signature was markedly enriched at six-week timepoint but not yet at 48 h after 6F transduction (**Fig. 2b**), and the CA activity measured from iPEC at six weeks was higher than that measured at four weeks (**Fig. 2c**).

**Figure 2.**
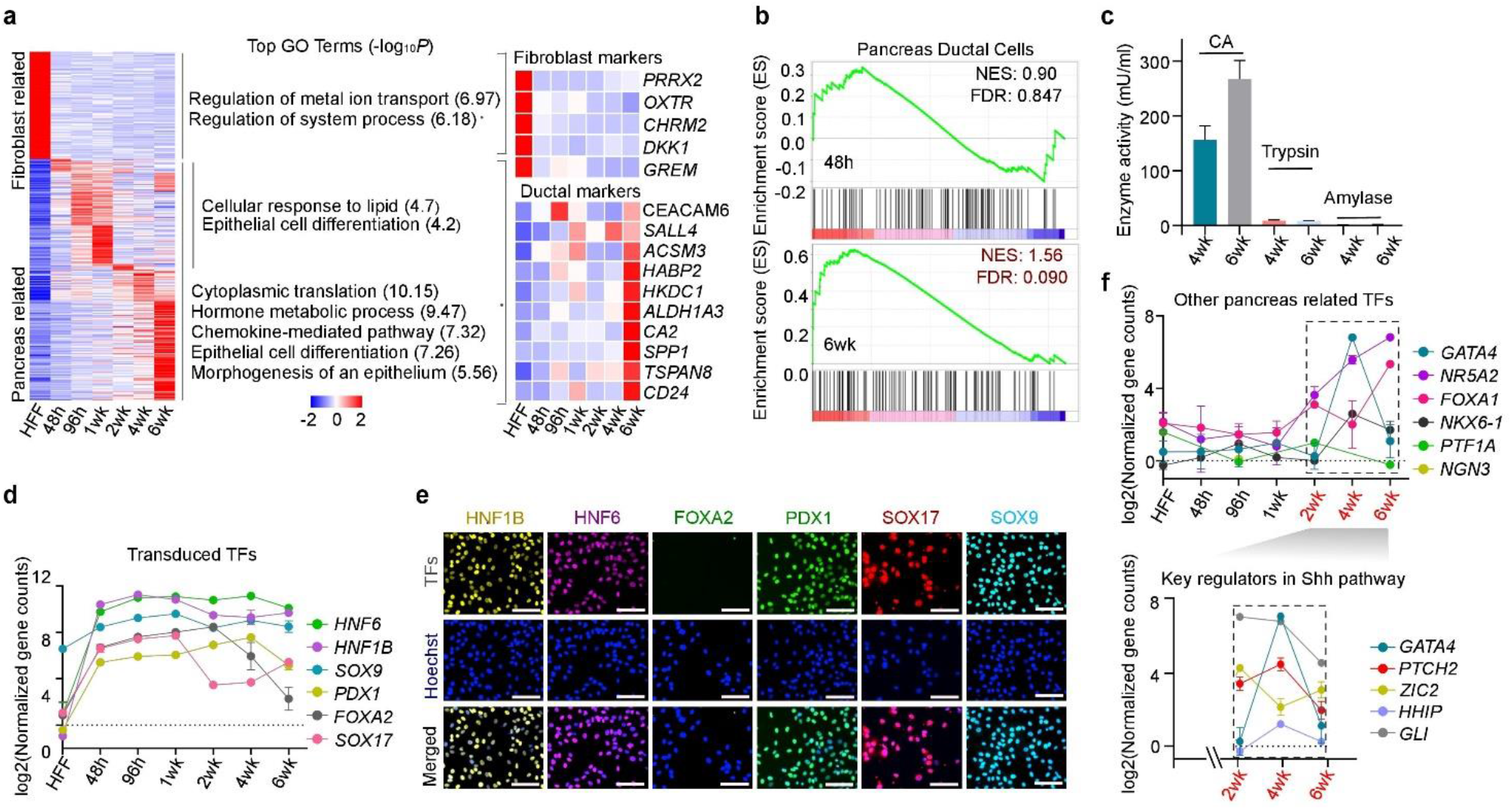
Gene expression dynamics during reprogramming. **a**, Heatmap clustering for average normalized expression values for DEGs (|log2FC| ≥ 1.5, *p*-value < 0.05) between reprogrammed cells at indicated time points and control HFFs; DEGs detected in all timepoints are shown. GO enrichment analysis for biological processes was performed for the indicated clusters using Metascape (Zhou *et al*, 2019). **b**, GSEA for enrichment of fetal pancreatic ductal cell signature in reprogrammed cells at 48 h and six-week timepoints. **c**, Measurement of enzyme activities for CA, trypsin and amylase in the lysates of iPECs at four- and six-week timepoints. Data are represented as the mean ± SEM (n = 6). **d**, Line plots showing the gene expression dynamics of 6F. **e**, IF staining of the TFs from 6F pool at six weeks after induction. **f**, Line plots showing the gene expression dynamics of other pancreas-related TFs. The normalized gene counts from RNA-seq data were log2 transformed and used for plotting in (**d**) and (**f**).

To further understand the dynamic control of cell fate conversion, we analyzed whether the TFs from the 6F pool are expressed consistently or show different patterns during reprogramming. Interestingly, the expression levels were largely consistent for *HNF6, HNF1B, SOX9* and *PDX1*, whereas *SOX17* and *FOXA2* were strongly downregulated around one to two weeks of reprogramming (**Fig. 2d**) despite being expressed from a constitutively active promoter. This is consistent with the earlier reported role of FOXA2 and SOX17 in specification of definitive endoderm in mice (Burtscher *et al*, 2009, Viotti *et al*, 2014) and role of FOXA2 in pancreatic differentiation from human pluripotent stem cells (Lee *et al*, 2019). At later stages from around four weeks onwards, the expression of SOX17 increased again but FOXA2 remained low, as seen from mRNA expression (**Fig. 2d**) and protein levels from IF staining (**Fig. 2e**). However, the expression of endogenous *FOXA1* increased, potentially compensating for the FOXA2 loss (**Fig. 2f**, *top*). These results suggest that SOX17 and FOXA proteins play distinct roles during the early and late stages of cell fate conversion.

For comprehensive understanding of the reprogramming process, we also analyzed expression patterns of endogenous pancreas-related TFs that were not part of the 6F reprogramming pool. This revealed striking temporal control of GATA4 expression with strong induction of its mRNA at four weeks and downregulation around six weeks of transdifferentiation, concomitant with sharp increase in FOXA1 expression (**Fig. 2f**, *top*). This suggests that GATA4 plays a critical role in establishing the pancreatic identity from early pancreatic progenitors. Previous studies have shown that GATA4 loss results in conversion of pancreatic cells to alternate intestinal and gastric cell fates due to aberrant activation of sonic hedgehog (Shh) pathway in pancreatic progenitors (Xuan *et al*, 2016) and that elevated levels of Shh signaling block pancreas formation (Hebrok *et al*, 1998). To identify the molecular determinants controlling Shh signaling during this critical step of pancreatic cell fate conversion, we analyzed the expression of canonical Shh signaling pathway genes from the Molecular Signatures Database (MSigDB; see **Methods**) in the RNA-seq data at two, four and six weeks of reprogramming. The highly stage-specific temporal activation of *GATA4* at four weeks was concordant with downregulation of *ZIC2* and upregulation of *PTCH2*, suggesting their role in suppression of Shh pathway that is required for pancreatic progenitor cell identity at this stage before ductal cell maturation **(Fig. 2f**, *bottom*). ZIC2, a downstream target of GATA4 (Rouillard *et al*, 2016), has been earlier reported to enhance Shh signaling through nuclear retention of Gli1(Chan *et al*, 2011) and PTCH2 is a negative regulator of Shh signaling (Alfaro *et al*, 2014). During later stages of reprogramming after pancreatic endoderm progenitor specification, the cells acquire ductal cell identity and the expression of GATA4 is downregulated (**Fig. 2f**, *bottom*). This is in agreement with previously reported GATA4 expression in the pancreatic progenitor state that is subsequently restricted to mature acinar cells during pancreatic maturation (Villamayor *et al*, 2020). Thus, our results give a mechanistic insight into how pancreatic identity is established and maintained through specific temporal suppression of Shh signaling.

Activation of other pancreas-related TFs such as *NR5A2* and *NKX6*.*1* was also observed at four weeks of reprogramming (**Fig. 2f**). *NR5A2* expression remained strong also at six weeks whereas *NKX6*.*1* expression decreased, commensurate with previously reported role of NR5A2 in pancreatic exocrine cells(Hale *et al*, 2014) and NKX6.1 expression restricted to β-cells in the mature pancreas (Petersen *et al*, 2018). On the other hand, TFs related to acinar cell specification such as *PTF1A* and endocrine cell fate such as *NEUROG3* were not detected during the reprogramming (**Fig. 2f**, *top*), indicating that the 6F pool mediates reprogramming of iPECs specifically towards pancreatic ductal cell identity.

### Global chromatin remodeling during iPEC reprogramming

To delineate the changes in chromatin state associated to TF-mediated lineage conversion, we performed ATAC-seq at the same timepoints that were used for RNA-seq within the first week of reprogramming. In agreement with the temporal changes detected in gene expression patterns, changes in chromatin accessibility were observed along the reprogramming time course and six broad clusters were identified (**Fig. 3a**). Based on ChIP-seq analysis for histone 3 lysine 27 acetylation (H3K27ac) at the same timepoints, the epigenome reprogramming through differential chromatin accessibility corresponds to changes in the levels of activating chromatin mark at the same loci (**Supplementary Fig. 2f**). Motif enrichment analysis of ATAC-seq peaks in different clusters revealed a shift in motif accessibility already around 96 hours of reprogramming from TFs that control the somatic cell identity of fibroblasts (FOS/TEAD/RUNX) to pancreas-related TFs corresponding to 6F pool such as HNF6, HNF1, SOX and FOXA (**Fig. 3a**). The same pattern was observed from differential motif analysis of accessible chromatin regions between one-week reprogrammed cells and HFF (**Fig. 3b**).

**Figure 3.**
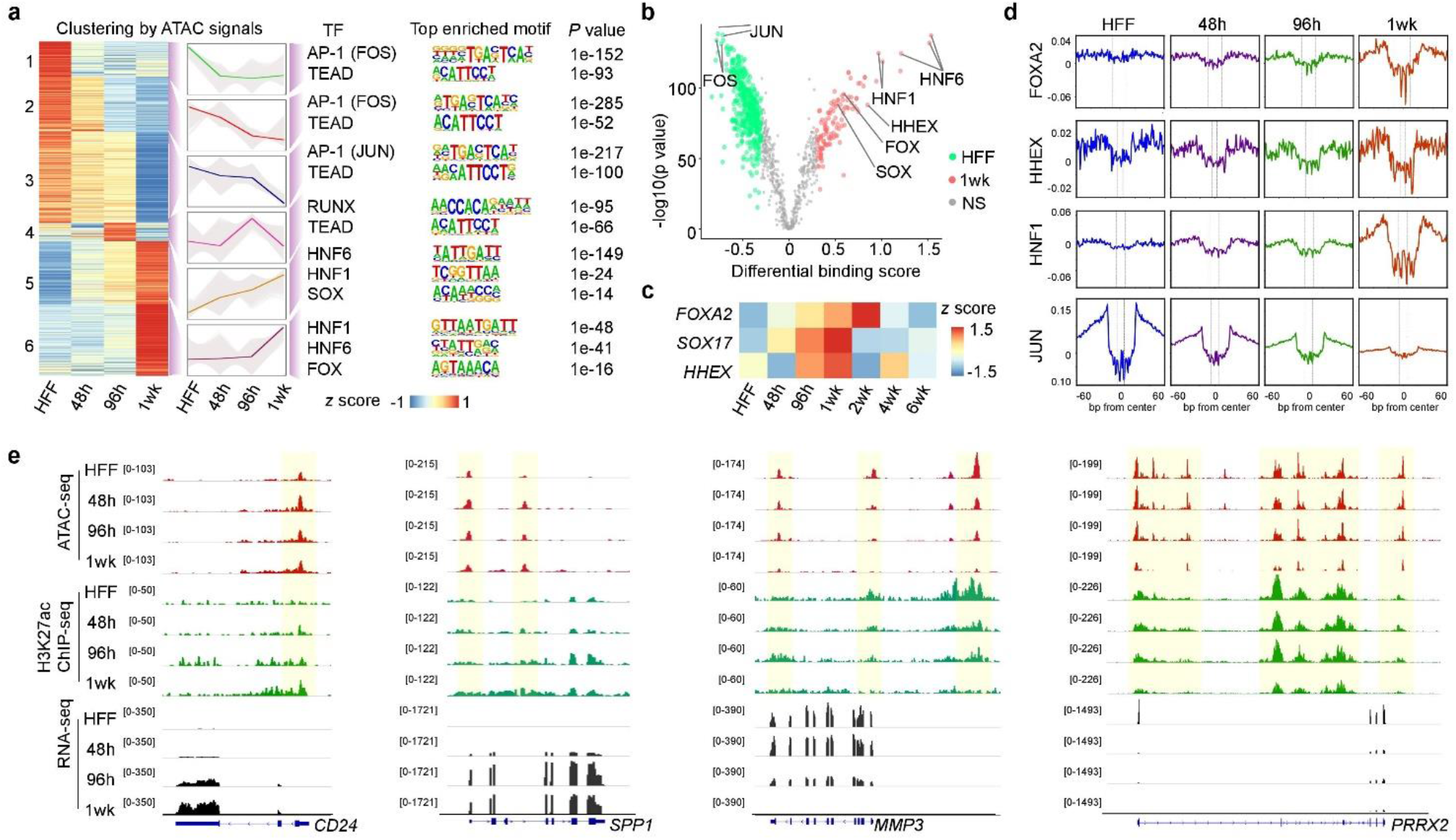
Chromatin accessibility dynamics during reprogramming. **a**, Unsupervised clustering heatmap of ATAC-seq data for peaks that show temporal dynamics of chromatin state at indicated timepoints after 6F transduction. Line plots showing the trend of each cluster. Top representative motifs ranked by *p*-value are shown. **b**, Volcano plot showing the differential predicted binding activity of all investigated TF motifs between HFF (left) and reprogrammed cells (right) at one week of reprogramming. TFs with |(differential binding score)| > 0.3 are colored on both sides. From each group, the top 1000 highest ranking peak regions according to RPKM values were used. **c**, Heatmap showing the z-score of normalized expression values for *FOXA2, SOX17* and *HHEX* from RNA-seq data at different timepoints during reprogramming. **d**, Aggregate accessibility profile of TF binding sites for selected differentially bound TFs shown in (**b**) from bias-corrected ATAC-seq footprints. Active binding of each TF is visible as depletion in the signal around the binding site. **e**, Genome browser snapshots showing representative marker genes of pancreatic exocrine cells (*CD24* and *SPP1*) and fibroblasts (*MMP3* and *PRRX2*).

In addition to the motifs for the 6F reprogramming TFs, a notable increase in the predicted binding activity was observed for HHEX between one-week reprogrammed cells and HFFs (**Fig. 3b and Supplementary Fig. 2g**). This was commensurate with a temporally controlled up-regulation of *HHEX* expression at 96 h and one-week timepoints (**Fig. 3c**). Interestingly, HHEX is known to be one of the earliest markers of the foregut progenitor cells that is required for formation of pancreas (Bort *et al*, 2004, Zhao *et al*, 2012) and for the specification of hepatopancreatic ductal system (Villamayor *et al*, 2020). Furthermore, TF footprinting analysis by TOBIAS framework (Bentsen *et al*, 2020) showed dynamic epigenome reprogramming of fibroblast chromatin during iPEC conversion with decrease in chromatin accessibility observed for TFs that maintain the somatic cell identity of the fibroblasts (JUNB, FOSL1) and increase for early endoderm specification TFs like SOX17, FOXA2, and HHEX, as well as pancreas-lineage determining TFs such as HNF6, HNF1B and SOX9 (**Fig. 3d and Supplementary Fig. 2h**). The changes in the chromatin state measured by ATAC-seq and H3K27ac ChIP-seq are commensurate with gene expression changes during reprogramming as seen from activation of pancreatic ductal marker genes such as *CD24, SPP1* and *SALL4*, and downregulation of fibroblast-related genes such as *MMP3* and *PRRX2* (**Fig. 3e and Supplementary Fig. 2i**).

Taken together, our data suggest that although cell fate conversion from one cell type to another using defined pool of TFs is a direct process, it involves transient intermediate progenitor states during which the cells move one step back towards a dedifferentiated state. This process is initiated by forced expression of strong lineage-dependent TFs such as FOXA2 and SOX17, leading to loss of the gatekeeper TFs (*e*.*g*. FOS/JUN/TEAD/RUNX) that maintain the fully differentiated somatic cell identities followed by endoderm specification through HHEX. Despite being transduced as a pool, the reprogramming TFs follow a distinct spatio-temporal activation and inactivation pattern, which implies that the direct TF-mediated cell fate conversion is controlled through three distinct steps: i) initiation of cell fate conversion by FOXA2 and SOX17, resulting in chromatin reprogramming, ii) cell fate specification and determination by HNF6 and HNF1B, and iii) cell fate maintenance and maturation by HNF6 and HNF1B together with SOX9 and pancreas-specific PDX1. This process also involves transient activation of distinct endogenous TF programs, namely HHEX at the endoderm progenitor state and GATA4 at the pancreatic progenitor state.

### Novel factor indexing approach for single-nuclei multiome sequencing (FI-snMultiome-seq) to dissect the role of defined TFs in gene expression and chromatin accessibility

To analyze the role of each individual TF and different combinations of TFs in cell fate control at a single-cell resolution, we developed a novel TF-indexing (barcoding) strategy, FI-snMultiome-seq, on top of the single-nuclei multiome sequencing platform from 10x Genomics for concomitant epigenome and gene expression profiling from the same nuclei/cell using snATAC-seq and snRNA-seq, respectively. For this, we designed a lentiviral expression construct harboring a unique sequence barcode at the 3’-UTR of the ORF (**Fig. 4a**). Each TF from the 6F pool was cloned into this expression vector, resulting in TF constructs with unique barcodes that are transcribed together with each TF and can thus be detected in the snRNA-seq experiment (**Fig. 4a**, *top*) after custom library preparation protocol. Importantly, the FI-snMultiome-seq approach is an improvement over the existing single-nuclei sequencing methods because it enables linking the TF expression directly to changes in gene expression and chromatin accessibility in each individual cell expressing one or multiple TFs, as is the case in reprogramming experiments. Moreover, the transcribing barcodes allow pooling of different experimental conditions into one Chromium multiome run, which is not possible with the currently available methods.

**Figure 4.**
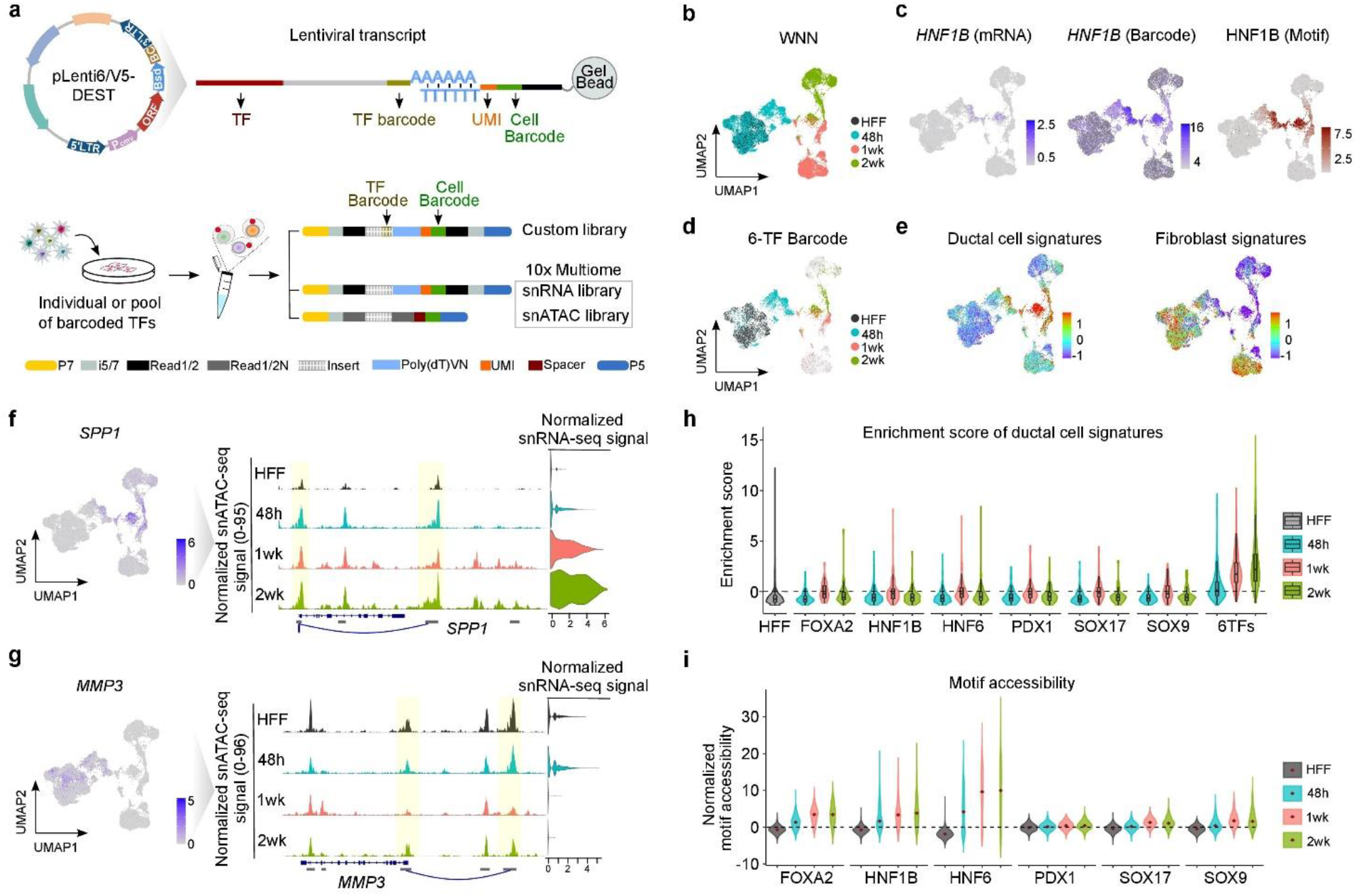
Novel FI-snMultiome method for dissecting the role of defined TFs in reprogramming at single-cell resolution. **a**, Schematic presentation of the lentiviral construct for ectopic expression of barcoded TFs, and the strategy for capturing the barcodes during 10x Multiome snRNA-seq workflow (top panel). UMI, unique molecular identifier. Reprogrammed cells transduced with six barcoded TFs individually or as a pool were harvested at different time points and multiplexed for the analysis of transcriptomic (snRNA-seq) and epigenetic (snATAC-seq) changes from the same cell. Custom TF barcode library is generated by an additional PCR after the pre-amplification step during the 10x Multiome workflow (see **Methods**). This enables correlating the TF barcodes with the 10x cell barcodes during the downstream analysis (bottom panel). **b**, Uniform manifold approximation and projection (UMAP) plot of all cells based on the integrated profiles of gene expression and chromatin accessibility using weighted nearest neighbor (WNN) analysis. **c**, UMAPs showing the expression of endogenous *HNF1B* mRNA (left), expression of transduced *HNF1B* detected from its unique indexing barcode (middle) and HNF1B motif accessibility analyzed from the snATAC-seq data (right). **d**, UMAPs highlighting the control cells and the reprogrammed cells expressing the barcodes for all six TFs. **e**, UMAPs showing the expression of ductal cell signatures (left) and fibroblast signatures (right) among all the cells. **f** and **g**, UMAPs and coverage plots showing chromatin accessibility at genomic loci of representative ductal cell marker *SPP1* (**f**) and fibroblast marker *MMP3* (**g**), along with the mRNA expression levels of the respective genes. **h**, Violin plot showing enrichment score of pancreatic ductal cell signatures in cells expressing individual TFs and the 6F pool and in HFFs as a control. **i**, Violin plot showing the motif accessibility scores (*z*-scores) of individual TFs in the cells with 6F pool at three timepoints and in control HFFs.

Barcoded TFs were transduced to HFFs either individually or as a 6F pool and the cells were maintained in the defined reprogramming media. The cells were harvested for FI-snMultiome-seq at three timepoints to analyze the intermediate stages through which individual cells progress during reprogramming (**Fig. 4a**, *bottom*). Sequencing library for the TF barcodes was prepared by an additional custom PCR step from the same pre-amplified material that is used for snRNA-seq and snATAC library preparation (see **Methods**). This results in three distinct libraries – snRNA-seq, snATAC-seq, and custom barcode libraries – that all are marked by the cell barcodes introduced by the 10x multiome workflow (**Fig. 4a**, *bottom*). Importantly, this strategy enables mapping each TF barcode to the cell barcodes and analyzing, which TFs are expressed in a cell and what are the corresponding chromatin and transcriptional states.

All cells harvested at different timepoints were analyzed together and projected into low-dimensional subspaces based on snRNA-seq for gene expression (**Supplementary Fig. 3a**), snATAC-seq for chromatin accessibility (**Supplementary Fig. 3b**), or the combined profile using both snATAC-seq and snRNA-seq measurements (**Fig. 4b**). The cells collected at 48 h are still closely similar to the control HFF, but the cells at one and two weeks after TF transduction have clearly distinct epigenome and transcriptome profiles (**Fig. 4b**). The FI-snMultiome-seq method has three major advantages: First, ectopically expressed transduced TFs can be robustly detected using the TF barcodes, giving high sensitivity for the analysis (**Fig. 4c and Supplementary Fig. 3c, d**). The motifs of 6F were also enriched within accessible chromatin in the cells where their barcodes were detected, suggesting the functional activity of the corresponding TFs in the transduced cells (**Fig. 4c and Supplementary Fig. 3e**). Second, current 10x multiome workflow is based on polyA capture and 3’ RNA-sequencing and is thus often unsuitable for capturing the transcripts originating from lentiviral expression vectors since the distance between the polyA (located in the 3’-LTR region) and the ORF of interest is usually long (>1 kb). In the FI-snMultiome-seq expression vector, the TF barcodes reside close to 3’-LTR (**Fig. 4a**), enabling their efficient capture and optimal library size for Illumina sequencing. This strategy also allows dissecting the effect of each transduced TF from the respective endogenously expressed TF based on the barcode and transcript (mRNA) expression, respectively (**Fig. 4c and Supplementary Fig. 3c, d**). Third, our approach enables analyzing only the cells that have been successfully transduced with TFs of interest and discarding all non-transduced cells from the analysis. Notably, the cell population that expresses all 6F barcodes (**Fig. 4d**) shows downregulation of the fibroblast signature and enrichment of the ductal cell signature (**Fig. 4e**). Further dissection of the signatures for different pancreatic cell types (Ianevski *et al*, 2022) revealed that reprogramming using the 6F pool results in clear enrichment of ductal cell marker genes whereas the signal for other pancreatic cell types such as acinar cells or endocrine cell types was weak (**Supplementary Fig. 4a**), confirming the ductal cell identity of the reprogrammed cells at single-cell level.

### FI-snMultiome-seq method reveals regulatory dynamics during cell fate conversion

The key feature of our FI-snMultiome-seq method is that it enables concomitant analysis of chromatin accessibility and gene expression from the same individual cells and specifically correlating them to TF expression. Different timepoints collected for FI-snMultiome-seq analysis (48 h, one week and two weeks after 6F transduction) allow temporal analysis of these regulatory changes during the early stages of reprogramming. Marker genes for ductal cells and fibroblasts, such as *SPP1* and *MMP3*, were clearly expressed in different cell populations (**Fig. 4f, g, Supplementary Fig. 4b**). Representative examples shown for the genomic loci of ductal cell marker genes (e.g., *SPP1, HNF6, HNF1B*) demonstrate dynamic reprogramming of chromatin accessibility already 48 h after 6F transduction commensurate with their increased transcript expression along the reprogramming time course (**Fig. 4f and Supplementary Fig. 4c**). In contrast, the chromatin accessibility was reduced at the loci for fibroblast-related genes (e.g., *MMP3, LOXL1, FBLN2*) in the cells expressing the 6F pool and their expression was downregulated or silenced at one-week timepoint (**Fig. 4g and Supplementary Fig. 4d**). These observations are in agreement with our bulk RNA-seq/ATAC-seq analyses that showed dynamic epigenome reprogramming from 48 h onwards, validating our population-level observations at the single-cell resolution.

Next, we used the FI-snMultiome-seq data to address whether all six TFs from the 6F pool are necessary for pancreatic ductal cell identity, since the factor-specific barcodes reveal the combination of TFs expressed in each individual cell. Previously, few reports have suggested that ectopic expression of a single TF can induce cell fate conversion (Chanda *et al*, 2014, Ng *et al*, 2021). For example, a pioneer factor *Ascl1* can efficiently convert fibroblasts to neurons (Chanda *et al*, 2014, Wapinski *et al*, 2013). Based on the FI-snMultiome-seq analysis, none of the individual TFs in the 6F pool was able to induce the expression of pancreatic ductal cell signature genes (**Fig. 4h**), suggesting that a concerted action of multiple TFs is necessary for efficient pancreatic exocrine cell fate conversion. Similarly, we analyzed the effect of all TF combinations from the 6F pool (five TFs, four TFs, three TFs and two TFs) on lineage conversion, showing that the 6F pool was the most efficient in induction of ductal cell signatures (**Supplementary Fig. 5a–d**). Furthermore, increased motif accessibility was observed for FOXA2, HNF1B, HNF6, SOX17 and SOX9 from 48 h to two-week timepoint compared to control fibroblasts (**Fig. 4i**). Taken together, 6F pool is the minimum combination of TFs for efficient specification of ductal-like lineage from human fibroblasts, and individual TFs contribute to the ductal cell fate in a sequential manner.

### Reconstructing the direct reprogramming path from fibroblasts to pancreatic ductal-like cells

To generate a high-resolution view of combined transcriptomic and epigenomic landscape in the heterogeneous cell population during transdifferentiation, we set out to reconstruct the reprogramming path based on pseudo-temporal ordering of the cells from the FI-snMultiome-seq data. A continuum of intermediate states through the first two weeks of reprogramming was captured as shown in **Fig. 5a**. This *in-silico* ordering of cells correlated well with the true time course of reprogramming (*c*.*f*. **Fig. 4d**). We next analyzed whether transduction with the 6F pool results in direct cell fate conversion from fibroblasts to iPEC or involves a transient intermediate step as suggested by our bulk RNA-seq and ATAC-seq data. Importantly, we detected specific expression of *HHEX* in the early stages of transdifferentiation (**Fig. 5b**) commensurate with a robust increase in the HHEX motif accessibility (**Fig. 5c**). This confirms our observation from the bulk analyses that showed specific increase in the *HHEX* expression and motif accessibility in 96 h and one-week reprogrammed cells *vs*. HFF together with other important pancreas lineage-specific TFs such as FOXA2, SOX17 and HNF1B (*c*.*f*. **Fig. 3b-d**). Importantly, upregulation of other definitive endoderm markers such as *KIT, KLF5*, and early endoderm regulon controlling TFs such as *YAP* (Moore-Scott *et al*, 2007, Nostro *et al*, 2011, Peng *et al*, 2020) coincides with the *HHEX* induction (**Supplementary. Fig. 6a**). Taken together, our results of iPEC conversion from HFF suggest that even though transdifferentiation using defined TFs is considered to be a direct cell fate conversion approach, cells make the transition through an intermediate progenitor state.

**Figure 5.**
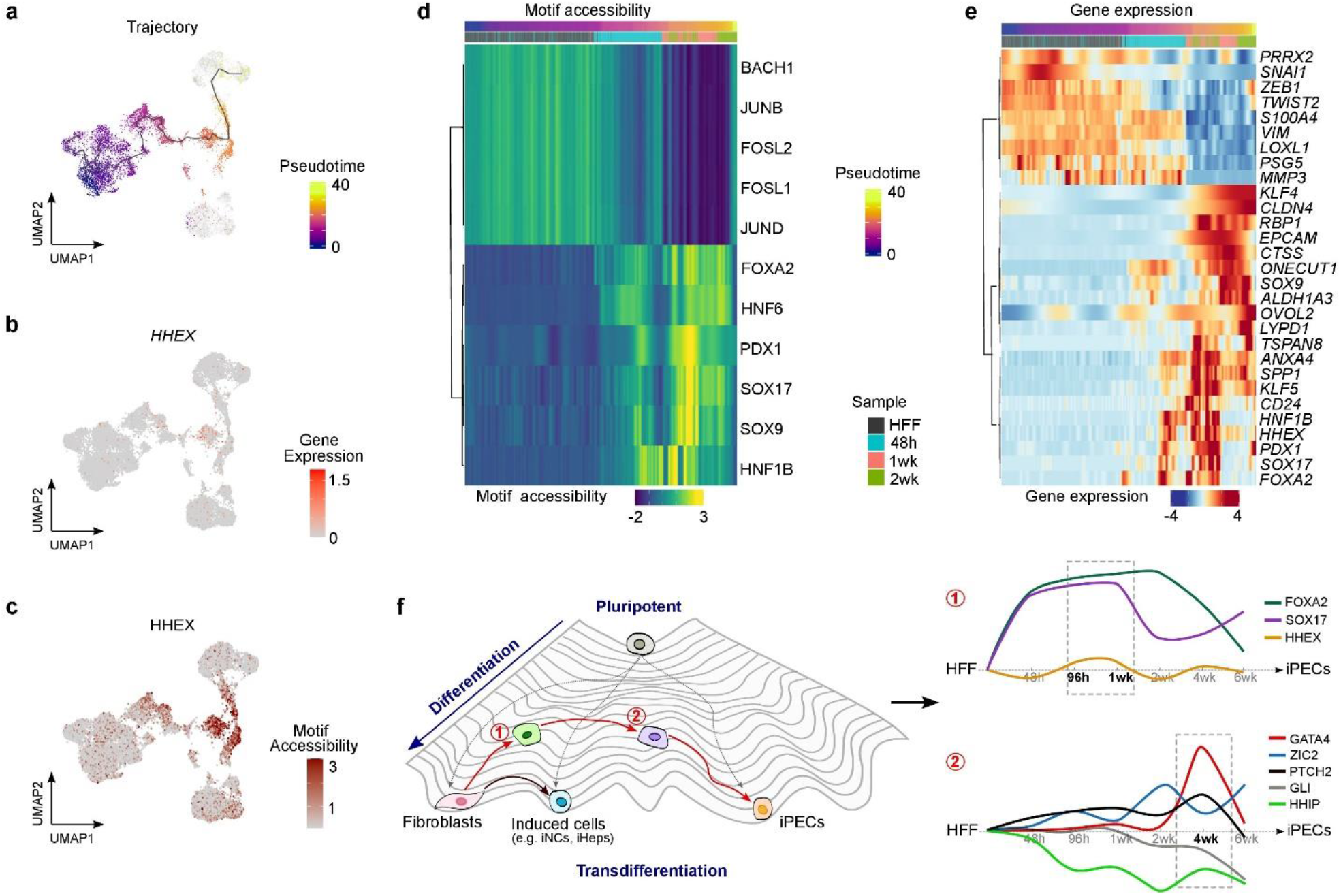
Reconstructing the reprogramming path from fibroblasts to pancreatic ductal-like cells. **a**, UMAP with cells colored by pseudotime. The gray line corresponds to the principal graph learned by Monocle 3(Cao *et al*, 2019). **b**, *HHEX* expression. **c**, Motif accessibility for HHEX. **d**, Heatmap showing smoothed pseudotime-dependent motif accessibility of representative motifs of somatic and pancreatic TFs. **e**, Heatmap showing smoothened expression levels of representative marker genes for fibroblasts and pancreatic cells. **f**, Schematic presentation of the transdifferentiation process from fibroblasts to iPECs in the context of Waddington’s epigenetic landscape, occurring through distinct molecularly controlled intermediate progenitor states: 1) definitive endoderm-like progenitor state controlled by FOXA2 and SOX17 and marked by HHEX expression, and 2) pancreatic progenitor state controlled by specific activation of GATA4.

Motif enrichment analysis along the pseudo-time course showed that motif accessibility increased markedly for pancreas-related TFs and decreased for fibroblast-related TFs (**Fig. 5d and Supplementary. Fig. 6b**). These pseudo-temporal changes also allowed to define the role of individual TFs in the 6F pool and to determine the regulatory events in the process of cell fate conversion. The first event at the earliest stages of reprogramming path is high FOXA2 motif accessibility (**Fig. 5d and Supplementary. Fig. 6b**), indicating its role in chromatin reprogramming and enhancer priming required for initiation of cell fate conversion. The second event in the iPEC reprogramming is increased motif accessibility for HNF6 (ONECUT1) that stays high towards the later stages as well (**Fig. 5d and Supplementary. Fig. 6b**), suggesting that HNF6 acts as the master TF for pancreatic exocrine cell fate specification. Third, the motif accessibility is increased for HNF1B, followed by the remaining TFs from the 6F pool – PDX1, SOX17, and SOX9, suggesting their role in pancreatic cell maturation and differentiation (**Fig. 5d and Supplementary. Fig. 6b**). Of note, the pseudotime analysis revealed a substantial increase in the motif accessibility for PDX1 that was not clearly visible from the analysis based on the sample timepoints only (*cf*. **Fig. 4i**), highlighting the strength of pseudo-temporal ordering in detecting the dynamic changes during reprogramming. As expected, motif accessibility for the TFs that maintain the fully differentiated somatic cell identity, such as FOSL1 and JUNB, decreased gradually along the pseudo-temporal axis (**Fig. 5d and Supplementary. Fig. 6b**). This chromatin accessibility shift is commensurate with changes in gene expression at the single-cell level since fibroblast-specific genes such as *VIM* and *MMP3* as well as TFs important for fibroblast identity such as *ZEB1, TWIST2*, and *SNAI1* were downregulated along the pseudotime (**Fig. 5e and Supplementary Fig. 6c**). On the other hand, expression of genes reported in early pancreatic progenitors such as *FOXA2, HHEX*, and *SOX17*, as well as pancreatic ductal cell and epithelial marker genes such as *SPP1, CD24*, and *EPCAM*, respectively increased along the *in-silico* pseudo-temporal ordering (**Fig. 5e and Supplementary Fig. 6c**), providing a time-resolved view of direct cell fate conversion from HFFs to pancreatic ductal epithelial cells.

Taken together, we have identified the minimum and sufficient combination of six TFs required for HFF conversion to iPECs of ductal epithelial cell type. Our results show that the transdifferentiation process is controlled by temporal and ordered activation of specific TFs that ensure cell fate establishment and maintenance, and that lineage conversion occurs through transient intermediate progenitor states, suggesting a paradigm shift in our understanding of transdifferentiation as a direct process in the context of Waddington’s epigenetic landscape (**Fig. 5f**).

## Discussion

Transdifferentiation is a powerful method for converting terminally differentiated mature cells to another specialized cell types through direct reprogramming without an intermediary pluripotent state. Although the TFs that are highly expressed in each cell type of interest are the likely candidates for the reprogramming factors, identifying the right combination of TFs that is necessary and sufficient for cell fate conversion has been challenging, owing to the enormous compendia of cell and tissue types in human body. Several groups have developed computational frameworks based on gene regulatory network analysis (Rackham *et al*, 2016, Cahan *et al*, 2014) to predict the combination of TFs required for cell fate conversion, but the *in silico* predictions are mostly for broad tissue types and not specific enough for distinct cell types. Thus, experimental approaches are critical for determining the specific combinations of reprogramming factors. Understanding the process of cellular differentiation and specification is particularly challenging when the target cell types arise from the same multipotent progenitors and express highly similar and related TFs, such as definitive endoderm-derived tissues pancreas, liver, and intestine. For instance, a combination of FOXA1 and HNF4A can give rise to liver- or intestine-like cells (Morris *et al*, 2014). For a more definitive hepatocyte identity, three TFs *i*.*e*., FOXA3, HNF4A and HNF1A are required (Huang *et al*, 2014), whereas addition of KLF4/5 to the pool of FOXA1 and HNF4A results in stronger intestinal-like identity(Morris *et al*, 2014). Here, we report the minimal and essential combination of six TFs, FOXA2, SOX17, PDX1, HNF1B, HNF6 and SOX9, that is sufficient to convert human fibroblasts to induced pancreatic exocrine cells with an epithelial-like morphology and global gene expression profile similar to pancreatic ductal cells. iPECs secrete carbonic anhydrase enzyme characteristic of functional ductal cells, indicating efficient cell fate conversion and maturation.

Lineage-specific TFs that control development and function of endodermal-origin tissues are largely similar, including HNF4A, forkhead family TFs FOXA1–3, homeobox family TFs HNF1A/B, and CUT domain TFs such as HNF6 (ONECUT1). Successful differentiation and maturation to any of these cell and tissue types requires coordinated action of specific TFs. During development, definitive endoderm is established by two TFs, FOXA2 and SOX17, and marked by HHEX. Expression of HHEX is critical to pancreatic development since *Hhex*^*-/-*^ mouse embryos were not able to specify a ventral pancreas (Bort *et al*, 2004). In our direct reprogramming model, FOXA2 and SOX17 expression together with other pancreas-related TFs in the 6F pool resulted in increased motif accessibility and expression of HHEX at one week after induction, observed from both bulk and single-nuclei ATAC-seq and RNA-seq data. Expression of other genes implicated in the endoderm such as *KIT, KLF5*, and *YAP* was also increased(Moore-Scott *et al*, 2007, Nostro *et al*, 2011, Peng *et al*, 2020). Interestingly, experimental and computational modelling has recently revealed the cell-fate plasticity of progenitor populations during pancreatic development (Willnow *et al*, 2021). Our results suggests that the direct lineage conversion occurs through an intermediate progenitor state that in the case of iPEC reprogramming is marked by HHEX positive cells. We propose that this is a transient touch-and-go step during which the cells dedifferentiate to progenitor endoderm-like state before committing to pancreatic identity.

Pioneer factors FOXA1 and FOXA2 have been implicated in the development of endodermal organs by priming organ-specific enhancers that trigger lineage-specific gene expression programs (Lee *et al*, 2019, Wang *et al*, 2015, Geusz *et al*, 2021). Current evidence from the developmental studies suggests that FOXA2 is the primary FOXA factor required for efficient pancreatic lineage induction (Lee *et al*, 2019, Geusz *et al*, 2021, Gao *et al*, 2008), but that they function in partially redundant manner (Geusz *et al*, 2021). Here, we discovered strong activity of FOXA2 motif at the early stages of reprogramming, indicating its role as a pioneer factor in chromatin reprogramming required for cell fate conversion. *FOXA2* expression was downregulated two weeks after 6F induction whereas the expression of *FOXA1* increased, potentially compensating for the *FOXA2* loss. Based on our FI-snMultiome-seq data, HNF6 was activated after FOXA2 but prior to the other factors such as HNF1B. In the mouse endoderm, Hnf1b is expressed before Hnf6 controlling its expression, but Hnf6 can also act upstream of Hnf1b (De Vas *et al*, 2015, Kropp *et al*, 2019, Poll *et al*, 2006). In our reprogramming system from human fibroblasts, both HNF6 and HNF1B were found to be necessary for cell fate conversion towards pancreatic identity. Furthermore, our results suggest that SOX17 has two distinct roles during conversion of fibroblasts to iPECs. First, it is strongly expressed at the endodermal progenitor state together with FOXA2 and HHEX but strongly downregulated at two-week timepoint. However, unlike FOXA2, SOX17 expression is induced again at six-week timepoint, suggesting that it contributes to the later stages of transdifferentiation as well, indicating its role in controlling two distinct programs. Collectively, the dynamic expression and chromatin accessibility of the reprogramming factors during iPEC conversion suggests that – although transduced to the cells as a pool and driven by a constitutively active promoter – these factors mediate the cell fate switch in a highly ordered and temporal fashion. Interestingly, sequential activation of pluripotency genes during induced pluripotent stem cell reprogramming has been shown to be controlled by dynamic reorganization of genome topology (Stadhouders *et al*, 2018), suggesting a plausible mechanism for the dynamic regulation of reprogramming factors.

Dynamic expression of many endogenous TFs, such as *GATA4, NKX6*.*1*, and *NR5A2*, was also observed during iPEC reprogramming. Interestingly, these factors were among the 14 candidate TFs for generating iPECs, but the reprogramming experiments showed that their ectopic expression was not necessary for establishing pancreatic ductal identity due to the activation of respective endogenous genes at specific timepoints during iPEC reprogramming. NR5A2 is an important regulator of pancreatic exocrine cell identity (Hale *et al*, 2014), and it was upregulated from one week onwards in the cells transduced with 6F pool. On the other hand, the expression of *GATA4* and *NKX6*.*1* was highly stage-specific, showing dramatic upregulation at four-week and downregulation at six-week timepoint. Previous studies have shown that GATA4 is required for pancreatic cell fate maintenance during pancreas development (Villamayor *et al*, 2020) but not for pancreatic cell fate specification since the mice with pancreas-specific *Gata4* deletion develop a pancreas (Xuan *et al*, 2012). Importantly, GATA4 is essential for maintenance of pancreatic progenitor cell fate by downregulation of Shh signaling (Xuan *et al*, 2016). In our reprogramming model, GATA4 upregulation was commensurate with downregulation of *ZIC2* and upregulation of *PTCH2*, resulting in negative regulation of Shh pathway since ZIC2 has been reported to activate (Chan *et al*, 2011) and PTCH2 suppress Shh signaling (Alfaro *et al*, 2014). In mature pancreas, *GATA4* expression is limited to acinar cells (Villamayor *et al*, 2020) and *NKX6*.*1* to β-cells (Petersen *et al*, 2018), indicating that their downregulation observed during iPEC reprogramming is necessary for the maturation of the ductal cell phenotype. These results provide detailed mechanistic understanding about the transdifferentiation process, highlighting the specific temporal activation of GATA4 that leads to downregulation of Shh pathway at the pancreatic progenitor stage that the cells acquire during reprogramming, and the requirement of GATA4 downregulation for the maturation of ductal cell identity.

Ectopic expression of lineage-specific TFs has become one of the most important approaches for cell fate conversion over the last two decades. However, not all cells in a typical reprogramming experiment can take or express the exogenous TFs regardless of their delivery method (viral/non-viral delivery), resulting in low conversion efficiency and mixed population of cells. Therefore, the key for efficient conversion and robust analysis is to identify the “real” reprogrammed cells from the heterogeneous population. Here, we developed a novel factor indexing strategy with transcribing barcodes on top of the Multiome platform from 10x Genomics. By tracking the TF barcodes, our FI-snMultiome-seq approach enables identifying the cells that express all stochastic combinations of transduced TFs, allowing not only to characterize the cells with different combination of TFs but also to evaluate the contribution of each individual TF to the acquired cell fate by comparing the cells expressing sub-pools of reprogramming TFs. In our FI-snMultiome-seq design, the TF barcodes reside only 78 bp upstream of polyA tail in the lentiviral constructs and are thus readily captured for the 3’ RNA sequencing utilized in the 10x Multiome workflow. Due to the longer distance between the ORF and the polyA region, the TF transcripts originating from the lentiviral constructs are not sequenced, and thus FI-snMultiome-seq approach enables segregation of transcripts originating from endogenous and exogenous factors by quantification of TF mRNA and barcode expression levels, respectively. This is particularly useful while studying the determinants of cell fate conversion, since the reprogramming TFs that are typically silent in the starting cell types might also get activated during the establishment of new cell fate. In conclusion, our FI-snMultiome-seq method provides a novel approach for studying the effects of distinct TFs on cell identity and their role in cell fate conversion.

Collectively, our results from conversion of fibroblasts to pancreatic ductal cells revealed that during direct reprogramming, the cells are highly heterogeneous and go through “chaotic” intermediate states marked by the expression of distinct progenitor markers. This is also seen from a gradual deactivation of TFs that maintain somatic cell identity such as AP1, TEAD and RUNX, implying that cells in this intermediate state have lost their original somatic phenotype and dedifferentiated back to a multipotent state. The right combination of TFs then activates the cell fate switch towards the right lineage through temporally controlled regulatory steps. These include i) cell fate initiation by FOXA2 and SOX17 along with activation of HHEX, mimicking the definitive endoderm progenitor state, ii) cell fate specification and determination controlled by HNF6 and HNF1B, involving a pancreatic progenitor state marked by GATA4 activation, and iii) cell fate maintenance and maturation controlled by HNF6, HNF1B, SOX9 and PDX1 that is associated to the downregulation of the progenitor-state signals. This coordinated process results in functional pancreatic ductal epithelial cells. Importantly, our novel protocol for generating pancreatic exocrine cells provides a platform for further studies of pancreatic malignancies as well as novel insights into pancreatic tissue engineering, suggesting a possibility of restoring impaired exocrine functions (Ellis *et al*, 2017). In the broader context of regenerative medicine, our paradigm shift findings about the transient progenitor states during direct cell reprogramming can help in identifying proliferative cellular states that are necessary for developing feasible cell-based therapies for various human disorders. In conclusion, our results provide a temporally resolved map of the molecular events that determine the defined cellular states during direct cell fate conversion, opening new avenues for understanding cell fate trajectories in light of these new findings.

## Acknowledgements

We thank Professor Lauri Aaltonen, Professor Jussi Taipale, and Dr Päivi Pihlajamaa for critical reading of the manuscript. We also thank HiLIFE research infrastructures including the FIMM Single-Cell Analytics unit, Biomedicum Flow Cytometry Unit, Biomedicum Imaging unit, Biomedicum Functional Genomics Unit (FuGU) and the FIMM NGS Genomics laboratory at the University of Helsinki. We thank the Center for Scientific Computing (CSC), Finland, for the computational infrastructure, Professor Lauri Aaltonen’s laboratory facilities for genomics work and Kerim Yavuz for technical assistance. This work was supported by the Academy of Finland (BS: 274555, 317807), Finnish Cancer Foundation (BS, SH), Sigrid Jusélius Foundation (BS, SH), Jane and Aatos Erkko Foundation (BS) and European Union’s Horizon 2020 Research and Innovation Programme under Grant agreement no. 965193 (DECIDER; SH).

## Author contributions

BS conceived and supervised the study, LF designed and performed all experiments, KZ analyzed the FI-snMultiome-seq data with the support from SH, and NP performed the bulk RNA-seq and ATAC-seq data analyses. BS and LF wrote the manuscript with contributions from all authors.

## Competing interests

Authors declare no competing interests.

## Additional information

Supplementary Information is available for this paper.

Correspondence and requests for materials should be addressed to Biswajyoti Sahu

## Data availability

All sequencing data generated in this study are available under GEO accession GSE216859. ENCODE blacklisted genomic regions for hg38 (accession ENCFF356LFX) were downloaded from ENCODE.

## Methods

### Cloning of human ORFs and lentivirus production

The full-length ORFs of all individual TFs were obtained from GenScript and cloned into lentiviral destination vector pLenti6/V5-DEST™ (Thermo Fisher Scientific, #V49610) using the Gateway™ LR Clonase™ II (Thermo Fisher Scientific, #11791020). The plasmid pLV-eGFP (#36083) was obtained from Addgene. For virus production, each TF expression plasmid was co-transfected with the packaging plasmids psPAX2 (Addgene #12260) and pMD2.G (Addgene, #12259) in 4:3:1 ratio into 293FT cells (Thermo Fisher Scientific, #R70007) with Lipofectamine 2000 (Thermo Fisher Scientific, #11668019). The culture medium was replenished on the following day and supernatant containing the viral particles was collected after 48 hours, filtered with 0.45 μm filters (Merck, #SE1M003M00), and concentrated using Lenti-X concentrator (Clontech, #631232), tittered using p24 assay and stored as single-use aliquots in −80 °C.

### Reprogramming protocol

Human foreskin fibroblasts (HFF; ATCC, #CRL-2429) were cultured in fibroblast medium (See **Supplementary Methods**). Early-passage HFFs were plated on matrigel-coated (Corning, #356230) dish on day 0 and transduced with constructs for TF expression (MOI = 1 for SOX17, FOXA2 and PDX1; MOI = 2 for HNF1B, HNF6 and SOX9) with 8 μg/ml polybrene on day 1. The medium was changed to fresh fibroblast medium on day 2. From day 3 onwards, the cells were cultured in defined reprogramming medium (See **Supplementary Methods**). During days 21–28, iPEC colonies were picked under microscope in a sterile hood, disassociated using Accutase (Gibco, #A1110501) and replated on matrigel-coated dish. Lentiviral expression construct for hTERT (MOI=0.5) was used for the immortalization of iPECs. The images of reprogramming cells were taken with ZEISS Axio Vert A1 microscope. All cell images throughout this manuscript were analyzed using Fiji (v1.53).

### Strategy for optimizing the combination of TFs for generating iPECs

In the first pilot experiment, HFFs were transduced with 13 TFs along with GFP (MOI=1; condition 1; **Supplementary Fig. 1b**) to induce pancreatic exocrine cell fate. The observed morphological and gene expression changes (*cf*. **Supplementary Fig. 1c**-**f**) suggested that some combination(s) of these candidate TFs can induce pancreatic identity, but the progress of cell fate conversion was slow and the number of cells showing morphological changes was small. We reasoned that constitutive expression of too many developmental TFs, or a potential lack of an essential TF may hamper efficient conversion. Therefore, we set out to optimize the reprogramming conditions, specifically by analyzing 1) if any TFs are redundant in the 13-TF pool, 2) if any essential TFs are still missing, and 3) by designing different reprogramming conditions for generating acinar- and ductal-like cells.

First, we excluded the two TFs (XBP1, GATA6) that were already expressed in HFFs and whose expression was not markedly induced at the two-week timepoint in the 13-TF sample. Based on the RNA-seq data, four of the TFs had basal expression in HFFs [mean log_2_(normalized gene count) from three replicates for XBP1, GATA6, SOX9, and HEYL being 10.9239, 7.4991, 6.5852, and 4.7049, respectively]. Further analysis of their expression in the 13-TF sample with mean log_2_(normalized gene count) of 11.2108, 9.2251, 9.0446, 10.8986, respectively, showed only weak induction for XBP1 and GATA6, suggesting their redundancy in the 13-TF pool. Moreover, GATA6 has been shown to be functionally redundant with GATA4 during pancreas development (Villamayor *et al*, 2020), further supporting its exclusion from the reprogramming pool. Next, we designed two sub-pools for generating acinar- and ductal-like cells separately based on previous information about human pancreas development (Jennings *et al*, 2015, Petersen *et al*, 2018), resulting in a 10-TF pool for acinar cells and a 9-TF pool for ductal cells (conditions 2 and 3, respectively; **Supplementary Fig. 1b**). For the ductal cell pool, we also included HNF1B that was not part of the initial 13-TF pool, due to its previously reported role in ductal cells of postnatal and adult pancreas (Solar *et al*, 2009). In total, 14 candidate TFs were included in the two reprogramming conditions.

After transducing HFFs with the TF pools for acinar and ductal cells (conditions 2 and 3), we observed reduced expression of fibroblast-related genes and gradually increasing expression of acinar and ductal cell markers, respectively, from qRT-PCR and RNA-seq analyses (*cf*. **Supplementary Fig. 1g**-**j**). In condition 2 for acinar cells, HFFs gradually lost their normal morphology after induction but we did not observe clear epithelial-like morphology during reprogramming. In condition 3 for ductal cells, some epithelial-like cells appeared at 21 days after induction, but these cells could not expand after passaging, potentially due to the small cell number. To increase the conversion efficiency, we further refined the TF pools on the basis of previous literature.

First, we excluded NKX6.1 and SOX9 from condition 2, resulting in an 8-TF pool for generating acinar-like cells (condition 5; **Supplementary Fig. 1b**). The exclusion was justified by the fact that both factors are expressed in pancreatic progenitor cells, but NKX6.1 is restricted to endocrine β cells during pancreas maturation (Petersen *et al*, 2018), whereas the cells that retain SOX9 expression undergo ductal fate specification (Shih *et al*, 2012). Furthermore, the cross-antagonism between Nkx6.1 and Ptf1a in pancreatic multipotent progenitors balance the endocrine and acinar cell neogenesis during normal development (Schaffer *et al*, 2010). However, in the reprogramming experiments using the condition 5 for acinar cells, we still did not observe clear epithelial morphology, and thus the experiments towards the acinar fate were not continued. Second, to refine the TF pool for the ductal fate, we excluded GATA4, NR5A2 and NKX6.1 from condition 3 based on their previously reported roles in pancreatic development, resulting in a 6F pool (condition 7; **Supplementary Fig. 1b**). In mature pancreas, GATA4 is largely exclusive to acinar cells (Villamayor *et al*, 2020) and NKX6.1 to β cells; NR5A2 is required for the development of exocrine pancreas but also plays an important role in acinar formation (Hale *et al*, 2014). Reprogramming experiments revealed that the 6F pool is able to generate pancreatic exocrine cells with phenotypic and functional characteristics of ductal epithelial cells (*cf*. **Figs 1, 2**). Other sub-combinations of candidate TFs were also tested, and the cells were characterized by bulk RNA-seq and/or FI-snMultiome-seq (*cf*. **Supplementary Fig. 1b, 5a-d**). Taken together, the combination of six TFs, FOXA2, SOX17, PDX1, HNF1B, HNF6 (ONECUT1) and SOX9, is necessary and sufficient for efficiently inducing pancreatic ductal cell fate from human fibroblasts.

### Immunofluorescence staining

Cells were fixed with 100% ice-cold methanol for 5 min at −20 °C and washed three times with PBS. After fixation, cells were permeabilized with 0.2% Triton X-100 for 15 min at room temperature (RT) and then washed three times with PBS. After permeabilization, cells were blocked by 5% BSA in PBS for 1 h at RT and then incubated with primary antibodies at 4 °C overnight. For the secondary antibody staining, cells were incubated with appropriate fluorescence-conjugated secondary antibody for 1 h at RT in dark. Nuclei were stained with Hoechst (Thermo Fisher Scientific, #62249). Primary and secondary antibodies were diluted in PBS containing 3% BSA. The antibodies used were : FOXA2, 20 μg/ml (R&D, #AF2400); SOX17, 15 μg/ml (R&D, #AF1924); PDX1, 1:50 (Santa Cruz, #sc-390792); HNF1B, 1:200 (Sigma, #HPA002083); ONECUT1, 1:50 (Santa Cruz, #SC-376167); SOX9, 1:1000 (Millipore, #AB5535); SPP1, 1:500 (Proteintech, #22952-1AP); E-cadherin, 1:50 (BD, #610182); donkey anti-goat (Alexa Fluor 488), 1:1000 (Abcam, #ab150129); goat anti-mouse (Alexa Fluor 488), 1:1000 (Thermo Fisher Scientific, #A11029); goat anti-rabbit (Alexa Fluor 594), 1:1000 (Thermo Fisher Scientific, #A11012); goat anti-rabbit (Alexa Fluor 488), 1:1000 (Thermo Fisher Scientific, #A11034). The images were taken with Nikon Eclipse Ti-E (with Primo) inverted microscope and analyzed using Fiji (v1.53).

### Flow cytometry analysis

Cells were harvested and centrifuged at 300g for 10 minutes. Cell pellet was resuspended in cell staining buffer containing 0.5% BSA and 2 mM EDTA. FcR blocking reagent (Miltenyi Biotec, #130-059-901) was used to reduce non-specific staining. Cells were then incubated with fluorescence-conjugated primary antibodies for 10 min in the dark at 4 °C. Cells were then washed twice in 1.5 mL of cell staining buffer and centrifuged at 300g for 10 minutes. Cell pellet was resuspended in 0.4 ml cell staining buffer for analysis. The antibodies used and their dilutions were: CD24-PE antibody, 1:20 (BioLegend, #311105); CD133-APC, 1:50 (Miltenyi Biotec, #130-113-746). Cells were sorted using BD Accuri C6 flow cytometer and the data were analyzed using FlowJo (v10.6.1).

### Enzyme activity analysis

Cells were harvested at different time points as indicated in the figures, washed with cold PBS and centrifuged for 3 minutes at 4 °C at 500g. Cell pellet was resuspended in 4X volumes of lysis buffer (150 mM NaCl, 10 mM Tris, 1% Triton X-100, pH = 7.2) and homogenized by vortex in cold room. Cell lysate was centrifuged for 2 minutes at 4 °C at 18,000g to remove insoluble material. Supernatants were then transferred to a clean tube and enzyme activities were analyzed using Carbonic Anhydrase Activity Assay Kit (Biovision, #K472), Amylase Activity Assay kit (Sigma, # MAK009) and Trypsin Activity Colorimetric Assay Kit (Sigma, #MAK290), following the instructions by the manufacturer. Optical density (OD) at 450 nm was read by FLUOstar Omega and the data were plotted using GraphPad Prism (v8.3.0).

### RNA isolation, qRT-PCR and bulk RNA-sequencing (RNA-seq)

iPECs were harvested for RNA isolation at 48 h, 96 h, one week, two weeks, four weeks, six weeks and 10 weeks after induction along with control HFFs. Total RNA was isolated using RNeasy Plus Mini Kit (Qiagen, #74134). For qRT-PCR, cDNA synthesis from at least three biological replicates was performed using the PrimeScript™ RT Master Mix (TaKaRa, #RR036A) and real-time PCR was performed using SYBR Green I Master (Roche, #04707516001). The primers used for each transcript are listed in **Supplementary Table 1**. The transcript levels of the target genes were normalized to GAPDH mRNA levels. Bulk RNA-seq libraries were prepared from 500 ng of total RNA for each sample using KAPA mRNA HyperPrep Kit for Illumina (Roche, #KR1352) following manufacturer’s instruction. Final libraries were quantified using Qubit HS Assay kit (Thermo Fisher Scientific, #15850210) and Tapestation High Sensitivity D5000 Reagents (Agilent, #5067-5593) for concentration and size distribution, respectively, then paired-end sequenced on Illumina Novaseq 6000.

### Chromatin immunoprecipitation-sequencing (ChIP-seq)

HFFs transduced only with GFP reporter construct and the iPECs at 48 h, 96 h and 1 week after 6F transduction were collected for ChIP-seq analyses. ChIP assays were performed as previously described (Sahu *et al*, 2011). Briefly, cells were fixed in 1% formaldehyde for 10 min at RT. Glycine was added to a final concentration of 0.125 M and incubated for 5 min to quench the reaction. Cells were then washed twice with ice-cold PBS and collected for lysis in RIPA buffer with protease inhibitors. Cross-linked chromatin was sonicated to an average fragment size of 200-500 bp, and immunoprecipitated with either H3K27ac antibody (Diagenode, #C15410196) or normal rabbit IgG (Santa Cruz, #sc-2027), 2 μg of antibody per reaction. ChIP libraries were prepared according to Illumina’s instructions and single-end sequenced on Illumina Novaseq 6000.

### Assay for Transposase-Accessible Chromatin-sequencing (ATAC-seq)

HFFs transduced only with GFP reporter construct and the iPECs at 48 h, 96 h and one week after 6F transduction were collected for analysis. ATAC-seq libraries were prepared from 75,000 cells as previously described (Buenrostro *et al*, 2015, Corces *et al*, 2017). Cells were washed with ice-cold PBS and resuspended in 50 µl of lysis buffer and incubated for 3 min on ice. The nuclei were isolated and transposed with Tn5 transposase in 2X tagmentation buffer (Illumina, #20034197) and incubated at 37 °C on thermomixer for 30 min at 1,000 rpm. The reaction was purified using MinElute PCR Purification Kit (Qiagen, #28004) and eluted in nuclease-free water. The samples were amplified for 8-11 total cycles and purified with AMPure beads (Agencourt, #A63881). Libraries were paired-end sequenced on Illumina Novaseq 6000.

### Factor barcoding design for FI-snMultiome-seq assay for snRNA-seq and snATAC-seq

For this assay, a 20-bp random oligo (N20) was introduced into the lentiviral expression vector pLenti6/V5-DEST™ by PCR (See **Supplementary Methods**). The PCR generates a unique indexing barcode for every single molecule and this unique barcode will be transcribed together with the respective ORF cloned into the expression vector allowing us to uniquely index and barcode each factor. For the barcoding PCR, 10 ng of pLenti6/V5-DEST™ vector was used as template per reaction with Q5® High-Fidelity DNA Polymerase (NEB, #M0491S). The PCR program, primers and the barcodes for each TF are as shown in **Supplementary Table 2**.

### FI-snMultiome-seq assay (snRNA-seq and snATAC-seq) from reprogramming cells

HFFs transduced only with GFP reporter construct and the iPECs at 48 h, one week and two weeks after induction were collected for analysis. Nuclei isolation for snRNA-seq and snATAC-seq was optimized and performed following the demonstrated protocol CG000365 from 10x Genomics. HFFs transduced with barcoded lentiviral expression constructs for 6F reprogramming TFs, both as individual TFs and as a pool of six TFs, were harvested at different time points after induction. For each time point, equal number of nuclei from each condition were pooled for library preparation. The libraries were prepared by the FIMM single-cell core facility (University of Helsinki) following the 10x Genomics user guide CG000338. An aliquot of the pre-amplified cDNA was used for preparing a custom library for detecting the TF barcodes. This strategy enables correlating each TF barcode to the cell barcodes introduced by the 10x multiome workflow. Custom barcode libraries were prepared using pre-amplified cDNA from Step 4 in protocol CG000338 by a two-step PCR process (See **Supplementary Methods**). The sequencing run parameters for the custom barcode libraries and GEX libraries were R1:28, i7-index: 10, i5-index: 10, R2:90 and for ATAC libraries R1:50, i7-index:8, i5-index:24, R2:49 on Illumina Novaseq 6000.

### RNA-seq data processing

The sample quality of the RNA-seq datasets were evaluated by FastQC analysis (http://www.bioinformatics.babraham.ac.uk/projects/fastqc/). Sequence reads were aligned to the human reference genome (UCSC GRCh38/hg38) using STAR aligner (v2.7.5c) with default parameters (Dobin *et al*, 2013). Samtools (v1.9) (Li *et al*, 2009) was used to sort the bam files. Gene counts were quantified using HTSeq-count (v0.11.2) (Anders *et al*, 2015). Genes with low counts having an expression of less than 10 across all samples were filtered out and the remaining genes were normalized using the DESeq2 (v1.30.1) pipeline(Love *et al*, 2014). To identify the differentially expressed genes (DEGs) between two conditions, a threshold of |log2FoldChange| > 1.5 and adjusted p-value < 0.05 was applied. Pathway enrichment analysis was conducted with a preranked gene list of all DEGs based on the sign of fold change and p-value using Gene Set Enrichment Analysis (GSEA) Preranked tool (v4.2.3). Principal component analysis (PCA) and volcano plots were generated using ggplot2 (v3.3.5) in R (v4.0.5). Heatmaps with hierarchical clustering using average linkage and euclidean as distance metric was created with pheatmap (v1.0.12). We overlapped the DEGs from all the time points in 6F pool to visualize the transient change in their expression pattern. The integrated average expression of the biological replicates, scaled and centered by row were plotted in the heatmap. Genes from the cluster were used to perform enrichment analysis for Gene Ontology terms by Metascape web-based platform (Zhou *et al*, 2019). To fetch the genes involved in Shh signaling pathway, we combined the gene sets of KEGG, Reactome, Pathway Interaction Database and WikiPathways from Molecular Signatures Database (MSigDB) and created a master set of 203 genes. We then analyzed the expression of these genes from our RNA-seq data at two, four and six weeks of reprogramming to generate a line plot.

### ATAC-seq and ChIP-seq data processing

ATAC-seq and ChIP-seq data processing was performed as previously described (Sahu *et al*, 2022). First, paired-end reads from all ATAC-seq timepoints were down-sampled to the similar sequencing read depth (36 and 32 million reads per sample for replicate 1 and 2, respectively). The quality metrics of the ATAC-seq and single-end reads from ChIP-seq fastq files were checked with FastQC. The reads were mapped to the human reference genome (UCSC GRCh38/hg38) by Bowtie2 (v2.2.5)(UCSC GRCh38/hg38) with option ‘--very-sensitive’ (Langmead *et al*, 2012). The ATAC-seq reads that mapped to mitochondrial DNA were discarded. Duplicates were marked using Picard MarkDuplicates (http://broadinstitute.github.io/picard/) and removed using samtools (v1.9) view with option ‘-F 1024’. ATAC-seq and ChIP-seq reads were subsequently filtered for alignment quality of less than 10 and 20 respectively. The peaks were called using MACS2 (2.1.1) with the parameter ‘--keep-dup all’ and the blacklisted regions downloaded from ENCODE were removed (Zhang *et al*, 2008). To generate the bigwig signal tracks from the aligned bam files, deeptools (3.1.3) bamCoverage was used with options ‘--normalizeUsing RPKM --binSize 10’ (Ramirez *et al*, 2014). Along with the biological replicates, we also generated i) pooled replicates by combining the reads of biological replicates and ii) pseudoreplicates which was made by randomly shuffling the pooled replicates and equal splitting. A similar peak calling strategy was used for these two methods as well and an overlap of peaks from all the methods was considered as the high confidence peak set for each time point. The peaks were further filtered using q value < 0.0001. The signal tracks from the pooled replicates were used for visualization in Integrative Genomics Viewer (IGV) genome browser (v2.5).

### ATAC-seq and ChIP-seq data clustering and motif searching

In order to observe the changes in chromatin accessibility across the time points, we employed CoBRA (v2.0) to pooled replicates of time-series data from ATAC-seq (Qiu *et al*, 2021). Briefly, the pipeline generates a master set of peaks from all samples and calculates the read density across those regions. Quantile normalization was used to normalize the read count matrix. The top 50% peaks were considered for the downstream unsupervised analysis and the rest were filtered out based on low reads per kilobase per million mapped reads (RPKM) values across multiple samples. k-means (k = 6) clustering was performed in the resulting peak set to highlight the substantial disparities in open chromatin profiles. Each cluster was paired with timepoint-specific line plots with a mean trend line using ggplot2 to visualize cluster trends. Peaks from each cluster were used to search for potential known and *de novo* motifs by using HOMER (v4.10.4) with the option ‘-size 100’ (Heinz *et al*, 2010). A similar approach was used for the H3K27ac ChIP-seq data to generate the master peak set. BEDTools (v2.30.0) intersect was implemented to investigate the overlap between H3K27ac peak set and ATAC-seq peaks from each of the clusters (Quinlan *et al*, 2010). The deeptools compute Matrix function was then used to generate a matrix of signal intensity around these overlapping peak centers (±2kb) of each cluster using the normalized bigwig files of H3K27ac ChIP-seq data. A heatmap was generated by plotHeatmap function using the matrix. Transcription factor Occupancy prediction By Investigation of ATAC-seq Signal (TOBIAS v0.13.3) was used to perform TF footprinting analysis in open chromatin regions(Bentsen *et al*, 2020). First, ATACorrect was used to perform bias correction of the reads in open chromatin by shifting +4 bp and −5 bp on positive and negative strands, respectively. Then we used ScoreBigwig to calculate footprint scores from the corrected cutsites with default parameters. Finally, we used BINDetect to find transcription factor motifs that were differentially footprinted at each time point using motifs from ‘The Human Transcription Factors’ (http://humantfs.ccbr.utoronto.ca/) database (Lambert *et al*, 2018). The aggregate footprint across the transcription factor binding sites of the differentially footprinted motifs were plotted using TOBIAS PlotAggregate.

### FI-snMultiome-seq (snRNA-seq and snATAC-seq) data processing

Raw sequencing data were processed using the Cell Ranger ARC pipeline (v2.0.1) with the GRCh38 reference (refdata-cellranger-arc-GRCh38-2020-A-2.0.0) for demultiplexing, alignment, barcode and feature counting to generate both ATAC and GEX feature-barcode matrices, which were loaded into Seurat (v4.1.1)(Hao *et al*, 2021) and Signac (v1.7.0)(Stuart *et al*, 2022) for further analyses. We filtered out cells with less than 1,000 RNA counts, 1,000 ATAC fragments or more than 100,000 RNA counts, 500,000 ATAC fragments, 30% mitochondrial-derived RNA counts. For scRNA-seq, the raw counts were normalized, scaled with the percentage of mitochondrial-derived counts and cell cycle scores regressed out using SCTransform. The top 3,000 highly variable features were used for principal component analysis (PCA), and the top 35 PCs were used for UMAP. For scATAC-seq, we called the peaks using MACS2 (v2.2.7.1). The peaks on nonstandard chromosomes and in GRCh38 genomic blacklist regions were removed. We performed the frequency inverse document frequency (TF-IDF) normalization using RunTFIDF, identified the top features using FindTopFeatures with min.cutoff = ‘q5’, performed latent semantic indexing (LSI) reduction using RunSVD, and the 2-35 dimensions used for calculating UMAP. We then constructed the WNN graph using FindMultiModalNeighbors with the 1-35 PCs from the scRNA-seq data and the 2:35 LSI dimensions from the scATAC-seq data for the joint UMAP visualization. Cells from different timepoints were analyzed in separate 10x Chromium runs but control HFFs transduced with GFP were pooled together with the 48h timepoint sample. During the analysis, the cells were negative to all TF barcodes were assigned as a control.

### snATAC-seq TF motif analysis

The same human motif position frequency matrices (Lambert *et al*, 2018) were used for snATAC-seq motif analysis that were used in the bulk analysis. Per-cell motif activity scores were computed using chromVAR (v1.18.0) (Schep *et al*, 2017) and the UCSC hg38 genome (BSgenome.Hsapiens.UCSC.hg38) with the Signac RunChromVAR wrapper.

### Cell type score

The cell type scores were computed using ScType (v1.0) (Ianevski *et al*, 2022) with cell-type-specific markers. The markers for pancreatic cell types were obtained from the ScType in-build database and the fibroblast markers were collected from literature (Muhl *et al*, 2020). All the markers used are listed in **Supplementary Table 3**.

### Trajectory and pseudotime analysis

The trajectory graph for control cells and the cells with six TFs was constructed using Monocle3 (v1.0.0) (Cao *et al*, 2019). The expression data and the joint UMAP projection was loaded into monocle3 using the as.cell_data_set function from SeuratWrappers (v0.3.0). The cells were clustered using the leiden method with the number of nearest neighbors set to 50, and the trajectory graph was learned using the learn_graph function with default parameters. Pseudotime was assigned using the order_cells function with setting the control cells as the starting points, and graph_test was used to identify genes and motifs that vary over pseudotime.

## Notes

### Competing Interest Statement

The authors have declared no competing interest.

